# Towards the Development of an Algae-Bacteria Pest Model: Enrichment of Oligoflexales Bacteria in *Nannochloropsis oceanica* Cultures with Different Starting Microbiomes

**DOI:** 10.1101/2025.04.17.649447

**Authors:** Thuy M. Nguyen, W. K. N. L. Abeykoon, Alina A. Corcoran

## Abstract

In 2018, a novel predatory bacterium named FD111 was identified, which could infect and significantly damage *Nannochloropsis* sp. in open pond cultures. Further research on this pest was hindered due to difficulties in cultivating it in laboratory conditions, making it tough to study its infection mechanisms and develop strategies for algal crop protection and Integrated Pest Management (IPM). After multiple attempts, a FD111-like bacterium pest was successfully resurrected using both a lab strain and a field-adapted strain of *Nannochloropsis oceanica*. The algae culture showed signs of deterioration after three days of infection, with a significant decrease in photosynthetic efficiency and bleaching of the culture. With the use of transmission electron microscope, a life cycle for FD111-like bacterium in *N. oceanica* was proposed, showing infection and reproduction mechanisms akin to earlier findings, albeit with slight differences in morphology. The FD111-like bacterium appeared to have a rod shape and flagella, which differed from the hook shape described in the only existing publication. Metagenomic sequencing identified the presence of three distinct sequences belonging to the *Pseudobacteriovorax* genus, within Oligoflexales order in the co-culture, suggesting multiple strains of FD111-like bacterium may be present. Future research will aim to obtain an isolate of the FD111 bacterium, gain deeper insights into its infection mechanisms, and develop protective strategies.

## 1. Introduction

*Nannochloropsis* is a genus of marine microalgae that has been extensively studied for its impressive lipid production, which can reach up to 70% of its dry weight depending on the environment in which it is grown [1,2]. The taxon has many industrial applications including the production of biodiesel, functional foods, and feed; wastewater treatment; and agriculture [3–5]. To meet commercial demands, *Nannochloropsis* is typically cultivated in open ponds, a cost-effective method that requires minimal maintenance [6–10]. However, this approach leaves the algae vulnerable to infections from predatory and antagonistic bacteria [11–16], resulting in a decrease in biomass yield during cultivation. For example, predatory bacteria like FD111 [17] have been observed attacking *N. salina* in outdoor algal ponds, causing significant damage to the cultures. These bacteria attached to *Nannochloropsis* cells, triggering clumping, discoloration, and eventual cell collapse. The transmission electron image illustrated the life cycle of the FD111 bacterium and its reproduction within host cell. The infection caused by FD111 was managed by supplementing with 2 mg/L of bleach, which prevented the culture from crashing. However, to date, there has been only one publication about this novel pest, the FD111 bacterium. There remains limited understanding of how FD111 bacterium infects *Nannochloropsis* sp. and how to protect algae from its attacks.

In another case, *Bacillus safensis* [18] demonstrated pathogenicity on *N. gaditana* when an additional 25% carbon source from marine broth media was provided, though no culture collapse was seen when the carbon source was omitted. This infection was controlled using a bacteriophage, a virus that preys on *B. safensis* cells while sparing the algal cells. Besides FD111 and *B. safensis*, *Vampirovibrio chlorellavorus* is a well-known pathogen of *Chlorella* species, particularly affecting *Chlorella sorokiniana* used in biofuel production [19–23]. As an obligate pathogen, it depends entirely on its host for survival and can cause significant biomass loss in algal cultures; however, successful isolation or replication of the *Vampirovibrio chlorellavorus* infection on *Chlorella sorokiniana* under laboratory conditions has never been achieved.

Bacterial infections remain one of the key factors dramatically diminishing the productivity of large-scale algae cultivation, and they are only three well-known pest models as mentioned previously. The scarcity of established laboratory models for studying algal-bacterial interactions hinders a comprehensive understanding of the mechanisms involved. Although numerous frozen samples of the FD111 infection were gathered from the field, efforts to revive this pest model in the lab had been unsuccessful for years. Consequently, we collected the infections once more from the crashed algal ponds, created a new batch of frozen samples, and proceeded with testing. This study aimed to successfully revive the FD111 pest from these frozen stocks. The research objective is to develop a reliable pest model that can be consistently reproduced in laboratory conditions, allowing us to study the infection process and propose possible defense strategies.

## 2. Materials and Methods

### 2.1. Strains and Cultivation Medium

To conduct our research, we use two different strains of *Nannochloropsis oceanica*. One strain, CCAP 849/10, was obtained from the Culture Collection of Algae and Protozoa (NCBI BioSample ID: SAMN39538505) in 2024. The other strain, called P7C12, was taken from an open pond at Sapphire Energy Inc., where it had been growing outside for over two years (NCBI BioSample Accession number: SAMN30478310). This strain was originally acquired as CCAP 849/10 in 2015 and maintained at Sapphire until inoculated outdoors. The field-adapted strain, P7C12, was suspected to be more resistant to bacterial infections, given its exposure to pests outdoors and/or evolution. Prior to and during experiments, both cultures were grown in 16NFL101 medium at a temperature of 27.5°C. They were exposed to continuous light at an intensity of 150 µmol/m^2^/s, provided by 4 bulbs 16W LED (Philips Lighting Inc., New Jersey, USA) shaken at 90 rpm on a VWR Advanced 5000 Digital orbital shaker (VWR International Inc., Pennsylvania, USA). The 16NFL101 medium contains the following ingredients per liter of Milli Q Deionized Water: 0.82 g sodium bicarbonate (NaHCO_3_), 16 g sodium chloride (NaCl), 0.59 g potassium chloride (KCl), 2.30 g magnesium sulfate heptahydrate (MgSO_4_*7H_2_O), 0.05 g calcium chloride (CaCl_2_) anhydrous, 0.354 mL urea ammonium nitrate solution 32 (UAN-32), 0.288 mL 8.5% phosphoric acid (H_3_PO_4_ v/v) – LC, 0.060 mL 100x CM Trace-Fe, and 0.024 mL of a 10000x Apollo Fe Stock, final pH of 8.0 [17]. The 100x CM Trace-Fe solution was prepared by dissolving 50 g of EDTA tetrasodium tetrahydrate salt (4Na-EDTA*4H_2_O), 4.58 g of manganese (II) chloride (MnCl_2_) anhydrous, 2.09 g of zinc chloride (ZnCl_2_) anhydrous, 1.26 g of sodium molybdate dihydrate (Na_2_MoO_4_*2H_2_O), and 0.40 g of cobalt (II) chloride hexahydrate (CoCl_2_*6H_2_O), then diluting the mixture to 1 L with Milli Q Deionized Water. The 10000x Apollo Fe Stock was made by combining 336.3 g of EDTA tetrasodium tetrahydrate salt (4Na-EDTA*4H_2_O) and 100 g of Ferix-3 and then topping off the solution to 1 L with Milli Q Deionized Water.

To infect the algae, we used two different sources of pests. The first source, known as NMP1, was collected from an open algal cultivation pond in New Mexico (NM) that had been infected by the FD111 bacteria and brought to the lab in May 2023. This source contained a mix of FD111 and other pests (e.g., ciliates, amoeba, flagellates). The second source, NMP6, was collected from mini ponds located at the Fabian Garcia Science Center during the same month. NMP6 was also a mixed pest source and underwent both passage and filtration through a 0.45 μm membrane to eliminate larger pests, retaining FD111 bacteria, which measured approximately 0.25 μm in diameter and 2-3 μm in length. We collected hundreds of 1.5 mL of crashed cultures from the field, stored them in 1.8 mL cryopreservation tubes, rapidly froze them in liquid nitrogen, and then kept them in a -80°C freezer until use. These samples remained in the freezer for about a year before commencing this experiment. Once removed from the freezer, the samples were left at room temperature in dim light to thaw before being used to infect the healthy algal cultures. We conducted qPCR on samples taken from algal cultures in an open pond, revealing that the FD111 bacterium was more prevalent than *Nannochloropsis* sp. and golden flagellates. This suggests that the FD111 bacterium was likely the main cause of the algal crash **(Fig. S3)**.

### 2.2. Enrichment of FD111-like Bacteria

We proceeded to transfer the FD111 infection source to a healthy *N. oceanica* culture, aiming to enrich and selectively cultivate the FD111-like bacteria over other bacteria present in the mixture. In 100 mL tissue culture flasks equipped with polypropylene screw caps vented by 0.2 µm membranes to increase air transmission and enhance cell growth (MTC Bio Inc., New Jersey, USA), 45 mL of algae at an exponential growth stage with an OD 750nm 0.1 was mixed with 5 mL of the infection source, resulting in a 10% v/v infection rate. The two strains of algae, CCAP 849/10 and P7C12, were infected with the two pest sources, NMP1 and NMP6, three replicates each treatment plus the algal control group. The cultures were maintained at 27.5°C, exposed to 150 µmol/m^2^/s light intensity, shaken at 90 rpm, and kept under continuous light. After being infected with bacteria for four days, the algal cultures began to deteriorate. The cells started to flocculate and changed from its original color to a brownish hue before eventually turning white as all of it died out. The infected cultures were then passaged to the healthy algal culture at an OD of 750nm, either 0.1 or 0.2. Each passage took 4 days, and this process was repeated ten times to increase the concentration of FD111 bacteria.

### 2.3. Metrics of Algal Biomass and Health

Each day, a 350 μm sample was transferred to a 96-well flat bottom black plate (Corning Inc., Maine, USA). The sample was then allowed to acclimate in the dark for 15 minutes. Using pulse amplitude modulation fluorescence (Mini-PAM 362 II, Walz, Germany), the photosynthetic yield of the dark-acclimated samples was measured. After measuring photosynthetic yield, a 100 µL sample was diluted at a ratio of 1:1 for OD 750 nm measurements and 1:20 for chlorophyll measurements. The OD 750 nm and chlorophyll fluorescence (430 nm absorbance, 685 nm emittance) were measured using a microplate reader (SpectraMax M2, Molecular Devices, USA).

### 2.4. Transmission Electron Microscopy

We utilized transmission electron microscopy (TEM) images to examine the life cycle and infection mechanism of the FD111-like bacterium. Additionally, we aimed to compare the morphology of the FD111-like bacterium observed in our current study with that documented in a previous publication. On days 0, 2, and 4, we collected 5 mL samples in 20 mL glass scintillation vials (Wheaton Inc., New Jersey, USA). Samples were fixed in 2.5% glutaraldehyde in 0.1 M sodium cacodylate buffer (pH 7.4) and stored at 4°C prior to processing. They were washed three times in 0.1 M sodium cacodylate buffer on a rotary mixer for 10 minutes per wash. Secondary fixation was performed with 2% osmium tetroxide in aqueous solution for 2 hours, followed by three 10-minute washes with deionized water. Dehydration was carried out through a graded ethanol series (30%, 50%, 70%, 90%, and 100%), with the final 100% ethanol step repeated twice. A 1:1 mixture of 100% ethanol and Spurr’s resin was prepared for sample infiltration for 1 hour, after which samples were transferred to 100% Spurr’s resin for an additional 1-hour infiltration. Samples were embedded in resin blocks and polymerized at 60°C for 48 hours. Embedded material was sectioned at 70 nm using a diamond knife (Diatome, Switzerland) on a Leica UC6 ultramicrotome (Leica, Germany). Sections were collected on 200-mesh copper grids, stained with 2% uranyl acetate for 20 minutes, washed with deionized water, counterstained with 3% lead citrate for 10 minutes, and washed again with deionized water. Sections were imaged at 80 kV on a Hitachi H7650 transmission electron microscope (Hitachi, Japan).

### 2.5. 16S Microbial Community Analysis

We employed 16S microbial community analysis to capture a snapshot of the bacterial composition during the algae culture crash. Additionally, we aimed to compare the microbial communities of the *N. oceanica* lab strain with the field-adapted strain. A 2 mL sample was snap frozen in liquid nitrogen and stored at a temperature of -80°C for subsequent DNA extraction. The samples were sent to the Microbiome Sequencing Service: 16S Amplicon Sequencing (Zymo Research, Irvine, CA) for processing and analysis. The DNA was extracted using the ZymoBIOMICS®-96 MagBead DNA Kit (Zymo Research, Irvine, CA) on an automated platform. Bacterial 16S ribosomal RNA gene sequencing was performed with the Quick-16S™ NGS Library Prep Kit (Zymo Research, Irvine, CA) targeting the V3-V4 region of the gene. This library preparation process involved real-time PCR reactions to control cycles and minimize PCR chimera formation. The resulting PCR products were quantified with qPCR fluorescence readings and pooled based on equal molarity before being cleaned and concentrated with the Select-a-Size DNA Clean & Concentrator™ (Zymo Research, Irvine, CA). Final quantification was done with the TapeStation® (Agilent Technologies, Santa Clara, CA) and Qubit® (Thermo Fisher Scientific, Waltham, WA) systems before sequencing on an Illumina® Nextseq™ with a P1 reagent kit (600 cycles). To improve sequencing accuracy, 30% PhiX spike-in was used during the sequencing process.

The raw data was processed using the DADA2 pipeline which included demultiplexing, joining pairs, and filtering for quality based on the q-score [24]. Next, the QIIME 2 2024.10 pipeline was applied to generate a table of features and select representative sequences [25]. Taxonomic analysis was performed using QIIME 2 with the SILVA 138.2 SSURef NR99 full-length database focusing on the V3-V4 region of 16S rRNA gene. QIIME 2 version 2024.10 was used for composition visualization, alpha-diversity, and beta-diversity analyses. Additional analyses, including heatmaps, Taxa2ASV Decomposer, and PCoA plots were carried out using PRIMER-e PERMANOVA+ version 7.

## 3. Results and Discussion

### 3.1. Enrichment of the FD111 Infection Source

In the first passage with NMP6, CCAP 849/10 and P7C12 both experienced crashes on day 4, whereas NMP1 decreased the growth rate of both algal strains, but didn’t cause a crash in either strain **(Fig. 1)**. By the 10th passage, both pests caused crashes in both algal strains on day 3. It was anticipated that continuous passages helped increase and selectively target the FD111 pest over other bacteria present in the samples **(Fig. 1)**, as well as remove any supportive bacterial communities in P7C12 strains. At passage 10^th^, the growth curve of the control group for both algal strains appeared similar, and the two infection sources also exhibited similar crashing patterns **(Fig. 1)**.

**Fig. 1.**
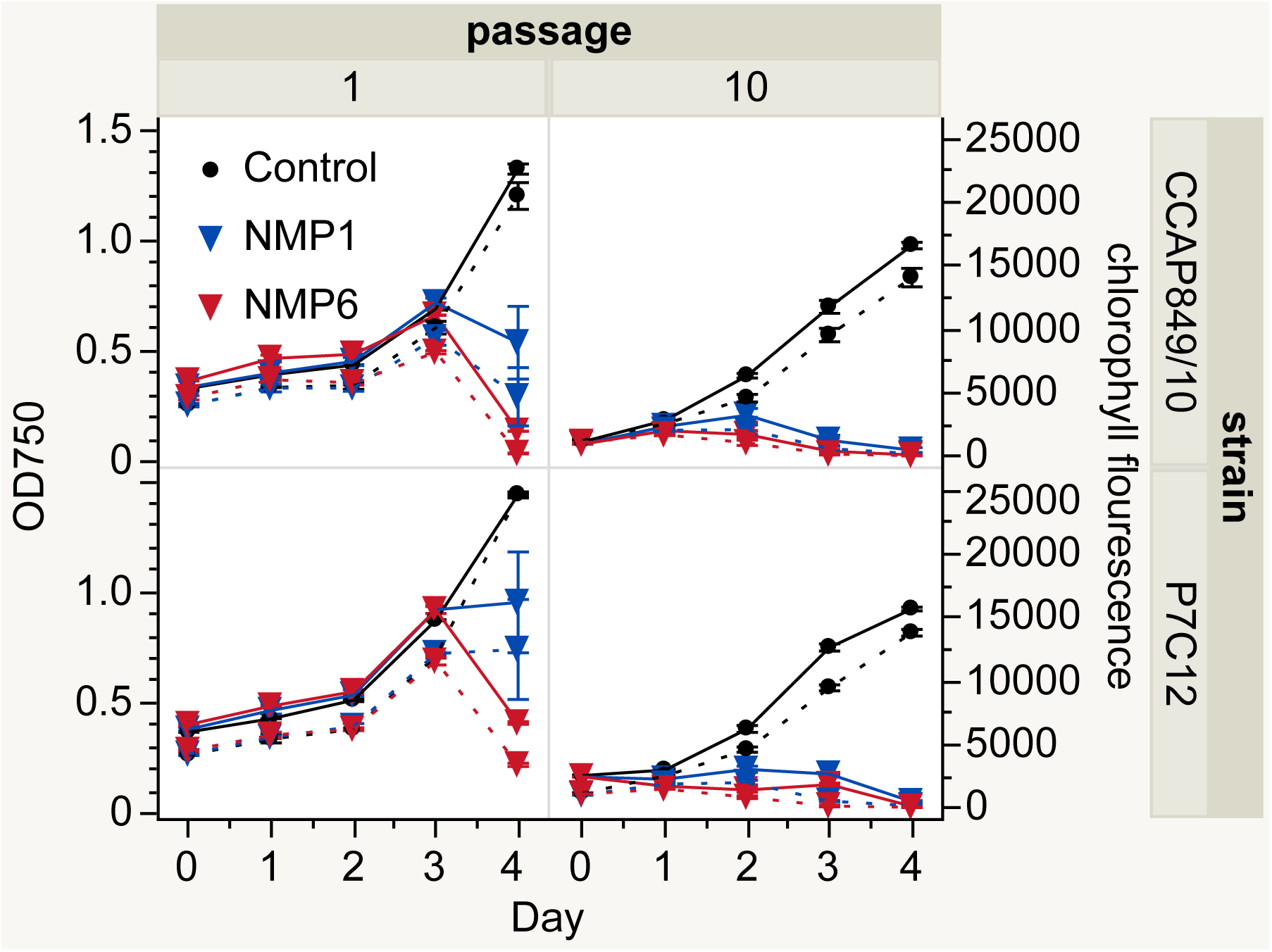
Optical density (left y-axis) and chlorophyll fluorescence (secondary right y-axis) of the *N. oceanica* lab strain (top, CCAP 849/10) and field-adapted strain (bottom, P7C12) crossed with two bacterial infection sources, NMP1 (non-filtered sample) and NMP6 (filtered sample). The solid line is OD750 value, while the dash line is chlorophyll fluorescence.

After four days, the OD750 reached 1 in the control group **(Fig. 1)**, as indicated by the greener color of the flask **(Fig. 2)**. In contrast, in the treatment group, both NMP1 and NMP6 doubled their biomass within one day of infection. By day four, OD750 and chlorophyll fluorescence dropped to 0, indicating complete death of the cultures **(Fig. 1)**. As a result of clumping together and dying on day three, the algae changed from green to yellowish brown and eventually turned white as they died off completely **(Fig. 2)**. Each passage experienced a crash after 4 days, and this consistent crashing occurred in more than 10 passages, with the crash behavior being successfully reproduced over a period of 1.5 months. However, the time it took for each crash to occur varied between passages **(Fig. S1).**

**Fig. 2.**
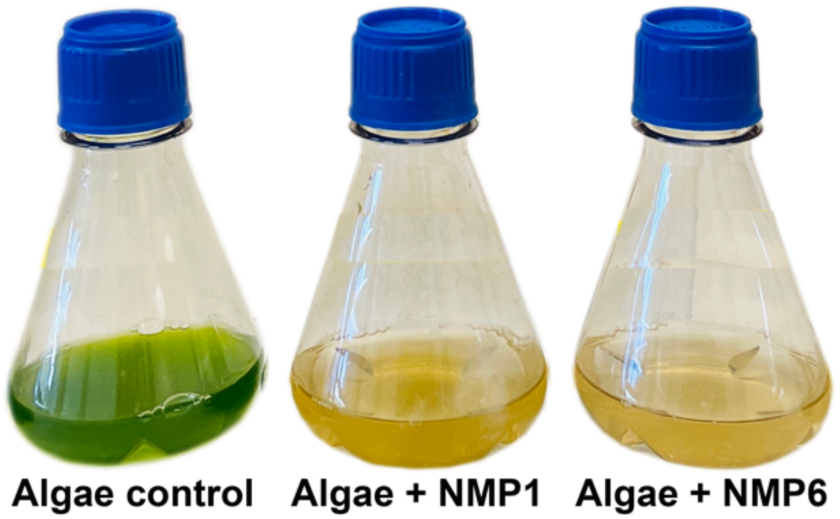
The algae in the control group remained green, whereas the algae with either NMP1 or NMP6 infection turned yellowish after four days of culturing.

### 3.2. Microscopy

The FD111 bacterium, discovered in 2018, was initially noted for its hook shape and the presence of a flagellum [17]. Upon the collapse of the algal culture, *N. oceanica* cells were seen clumping together under light microscope. The microscope image also revealed that FD111-like bacterium exhibited a rod shape **(Fig.3.A)**. Furthermore, TEM analysis indicated that the morphology of FD111-like bacterium has changed. Infections from both NMP1 and NMP6 samples now displayed rod-shaped with a flagellum **(Fig.3.B)**. Based on observations from both light microscopy and TEM, it was concluded that the FD111-like bacterium in this study possesses a rod shape.

**Fig. 3.**
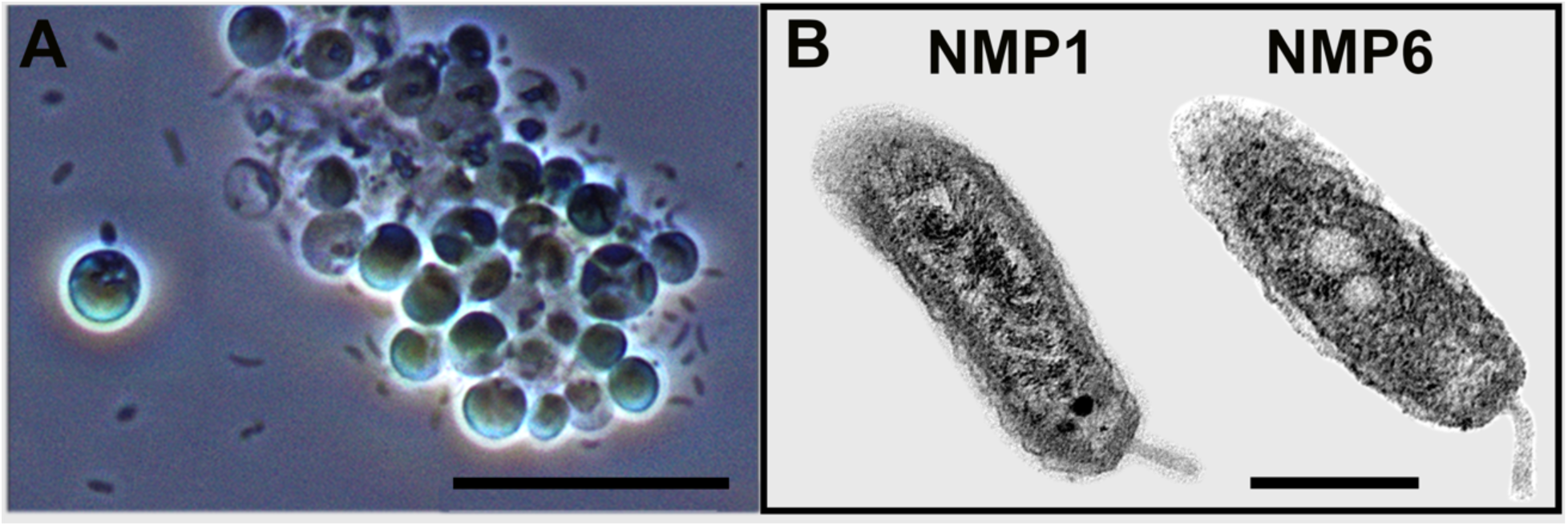
(A) Light microscope images of *N. oceanica* infected by FD111-like bacterium at 10 μm. (B) TEM images of FD111-like bacterium from NMP1 and NMP6 samples AT 500 nm.

**Fig. 4.**
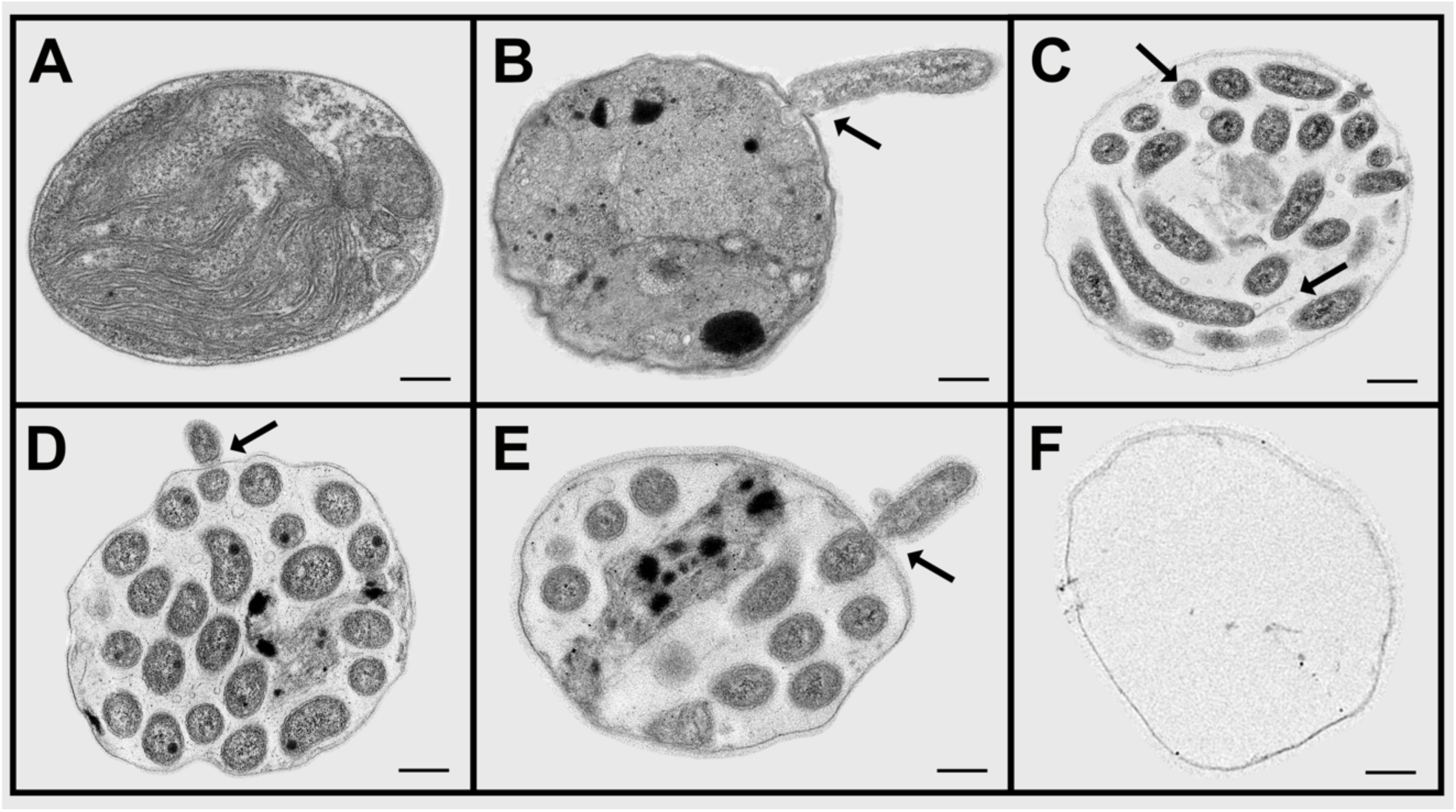
The potential life cycle of an FD111-like bacterium with a *N. oceanica* algal host as observed by TEM is as follows: **A.** Healthy *N. oceanica* algal cell alone. **B.** FD111-like bacterium attached to the algal cell wall and transferred its cellular material into the host cell through a pore in the wall. **C.** FD111-like bacteria with flagellum at longitudinal shape, and in a coiled shape. **D & E.** FD111-like bacterium exited the algal host cell. **F.** Algal cell wall is left empty, without any organelles. Scale bar: 200 nm.

TEM images showed the potential life cycle of FD111-like bacteria interacting with the *N. oceanica* algal host. (A) The healthy *N. oceanica* algal cells appear with intact organelles, a full chloroplast, and a complete cell wall. (B) The infection bacteria attached to the algal cell wall of *N. oceanica* algal and injected its reproductive materials through the pores in the algal cell wall (black arrow), causing the wall became wrinkled. (C) FD111-like bacteria reproduced inside the algae host cell, appearing in a longitudinal form with a flagellum and the coiled shape. All the algal organelles were consumed by infected bacteria. (D & E) FD111-like bacteria existed the host algal cells through the pore, leaving behind an empty algal cell wall (F). The infection and reproduction mechanism is similar to previous publication [17].

### 3.3. Microbial Community Structure

Prior to inoculation with FD111, there was a difference in the microbial communities between the *N. oceanica* lab strain CCAP 849/10 and the field-adapted strain P7C12 at day 0 **(Fig. 5)**. In CCAP 849/10, bacteria were more prevalent than algae, while in P7C12, algae were the most dominant. Over a 4-day culturing period, the microbial community of the control underwent changes. By day 4 in the control, the most dominant groups included *Nannochloropsis*, Rhodobacterales, Flavobacteriales, Rhizobiales, Sphingomonadales, and Cytophagales (**Fig. 5)**. At this point, when the algal culture completely collapsed, the microbiome profiles of the four treatment combinations appeared similar. The *Nannochloropsis* algae were entirely absent in the crashed cultures as predatory bacteria had entirely devoured them. The dominant microbiome also shifted with the culture crash, with Micrococales becoming the most dominant order in the treatment. Rhodobacterales, Balneolales, and Flavobacteriales were also observed as dominant groups in the crashed culture (**Fig. 5)**. The FD111-like bacteria from the Oligoflexales order were not found in the control for either algal strain at any time point. It began to appear in the treatment when FD111-like bacteria were introduced to the treatment at day 0, but it was not the most dominant organism in the co-culture when the algal culture crashed (**Fig. 5)**.

**Fig. 5.**
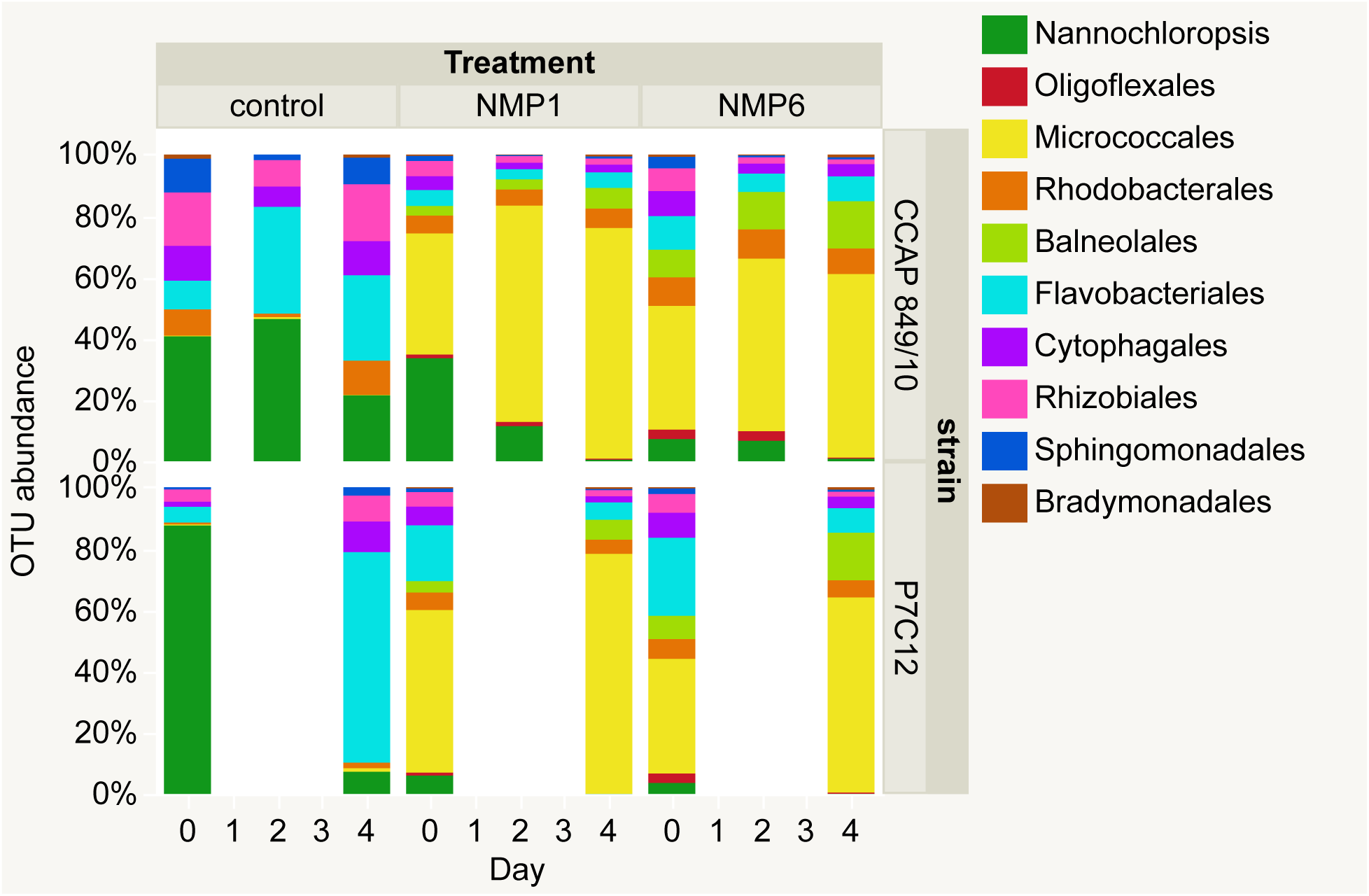
The OTU abundance of the top 10 Orders in the microbial community was analyzed for two *N. oceanica* strains after cross-infection from two sources over four days. Samples from the lab strain CCAP 849/10 were taken on days 0, 2, and 4, with duplicates, while samples from the field-adapted strain P7C12 were collected on days 0 and 4, without any replication.

The heatmap **(Fig. 6)** displayed the abundance of OTU in the *N. oceanica* lab strain, including both the algae control and those cross-infected with two sources over 4 days. The control samples, which consisted solely of algae, were grouped together, while the treatments were organized by time point. Micrococcales emerged as the most dominant order in the treatments. Four orders - Propionibacteriales, Saccharimonadales, Oligoflexales, and Sphingobacteriales - did not appear in the control but were present in the treatments and during the culture crashes. Oligoflexales was absent from the control at any time point but was found in all treatments, being most abundant in the NMP6 - filtered sample, replicate 1 on day 2, just before the culture crashed.

**Fig. 6.**
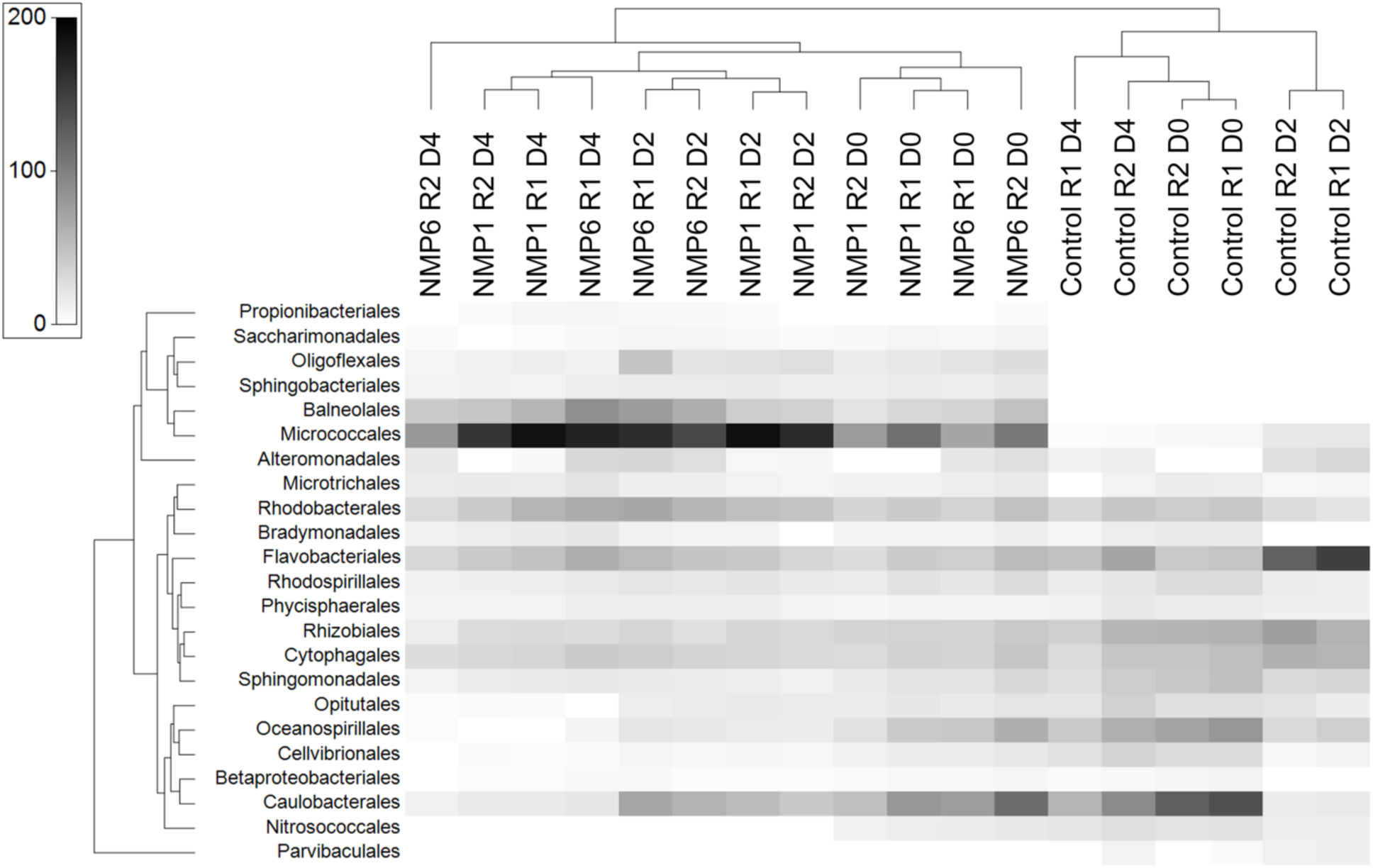
The heat map shows OTU abundance in *N. oceanica* lab strain, including the algae control and those cross-infected with two sources over 4 days. “R” stands for replicate, “D” stands for day.

The sequencing reads from the co-culture of *N. oceanica* and two infection sources were blasted again the SILVA 138.2 SSURef NR99 full-length database, specifically targeting the V3-V4 region of the 16S rRNA gene. This analysis identified three species within the Oligoflexaceae family, all classified under the *Pseudobacteriovorax* genus. The only species-level match found in the database was identified as *Pseudobacteriovorax antillogorgiicola*, a gram negative bacterium, within the order Bdellovibrionales [26,27]. It was collected and isolated in 2025 from the coast of San Salvador, The Bahamas. TEM morphology of *P. antillogorgiicola* as having a rod shape with a long curly tail [26]. However, the TEM in the current study showed differences, featuring a shorter tail and a more symmetrical body **(Fig. 3.B)**. This morphological difference suggests that the known *P. antillogorgiicola* and the FD111-like bacterium are distinct organisms. In this study, the other two *Pseudobacteriovorax* species found belonged to unculturable taxa, with one identified only at the genus level. It seems that there has been minimal research aimed at isolating these organisms and comprehending their physiology and infection mechanisms, leaving an opportunity for additional research to further explore this subject.

## 4. Conclusions and Future Scope

The FD111 infection samples were gathered from algal ponds that had been crashed by a bacterial invasion. Numerous cryopreserved vials were stored at -80°C for later research purposes. The infection was revived for two years at Sapphire Inc. and for one year at Los Alamos National Lab (LANL) as noted in the PEAK report [28]. However, it eventually became non-functional at LANL, the New Mexico Consortium, and New Mexico State University. After numerous failed attempts, we have finally managed to replicate the crash of FD111 – *N. oceanica* in a controlled laboratory setting. This is a significant milestone for future research as it allows us to further study the mechanisms of infection. For example, we can isolate FD111, manipulate the culture conditions to study its impact on the crash, conduct omics analysis to understand the infection process better, and evaluate other affordable methods for their effectiveness in controlling the crash.

NMP1, the non-filtered sample, did not lead to any crashes in either of the *Nannochloropsis sp*. strains during the first passage. However, it did cause both strains to crash by the 10th passage. The crashing pattern of NMP1 at the 10th passage resembles that of NMP6 as FD111-like bacteria (*Pseudobacteriovorax* genus) were present across all the crashed samples. Continuing to transfer the infection would increase the concentration of the pests, providing a method to maintain an active infection source within the lab culture environment and enrich it. In addition to continuous passages, other techniques like plaque agar plates or flow cytometry machines should be utilized to separate pest cells from algae cells.

The *N. oceanica* field-adapted strain P7C12 was maintained in an outdoor open pond at a private industrial company for 1.5 years. It was known to survive in the presence of the FD111 bacterium. Genomic sequencing revealed that the genome of the *N. oceanica* field-adapted strain remains similar to the lab strain CCAP 849/10 [29]. However, it was hypothesized that it had a more supportive microbial community and greater resilience to predatory bacteria. In this study, the P7C12 strain was crashed in the first passage by the NMP6 infection source, contradicting the original hypothesis. The 16S rRNA microbial community data also showed no significant differences in the bacterial community between P7C12 and CCAP 849/10.

In 2018, the FD111 pest was first discovered in a hook shape with a flagellum [17]. However, this study reveals that the pest now has a rod shape with a flagellum. There is no discernible difference in morphology between the two infection sources NMP1 and NMP6. FD111 bacteria were initially grouped under Bdellovibrio-and-like organisms due to their predatory nature [30],[17]. Their morphology shifts between a predatory and a non-predatory phase [31,32]. In the predatory stage, they exhibit a vibrioid shape with a polar flagellum, while during the non-predatory stage, their cellular shape varies significantly [31–33]. The morphological differences of the FD111-like bacteria observed in this study compared to earlier research might suggest several possibilities: it could represent new strains, and the intensity of its predatory characteristics may have evolved over time. Further shotgun sequencing is needed to fully understand the genetic information of the published FD111 genome compared to the genome of the pest in this study and determine if they are identical or distinct strains.

The 16S rRNA sequencing results identified three distinct bacteria from the *Pseudobacteriovorax* genus within the Oligoflexales order. It is likely that the mixture contains three different FD111-like variants. The data was examined using the SILVA 138.2 SSURef NR99 full-length database, focusing on the highly conserved V3-V4 region of the 16S rRNA gene. Introducing the mixture to various algae strains, which offer alternative food sources, could help enrich these FD111-like variants, leading to their isolation and subsequent evaluation of their individual effects.

This study contributed to expanding the existing knowledge about the FD111-like bacteria and algae-bacterial pest model. It detailed the laboratory setup and culturing conditions necessary to revive the FD111-like bacterium in a controlled environment. This significant contribution lays a solid groundwork for future research aimed at identifying other factors in the mixture that might lead to algal crashes and finding ways to reduce the impact of contaminating bacteria. Additionally, we aim to gain a deeper understanding of the infection mechanisms, which will help us propose effective, low-cost treatments to safeguard algal cultures.

## Funding

This work was funded by DOE BETO Grant EE-EE0011057 Antivirulence Approaches to Treat Algal Crops (AvATAC) to Alina A. Corcoran & John McGowen.

---

**Figure S1.**
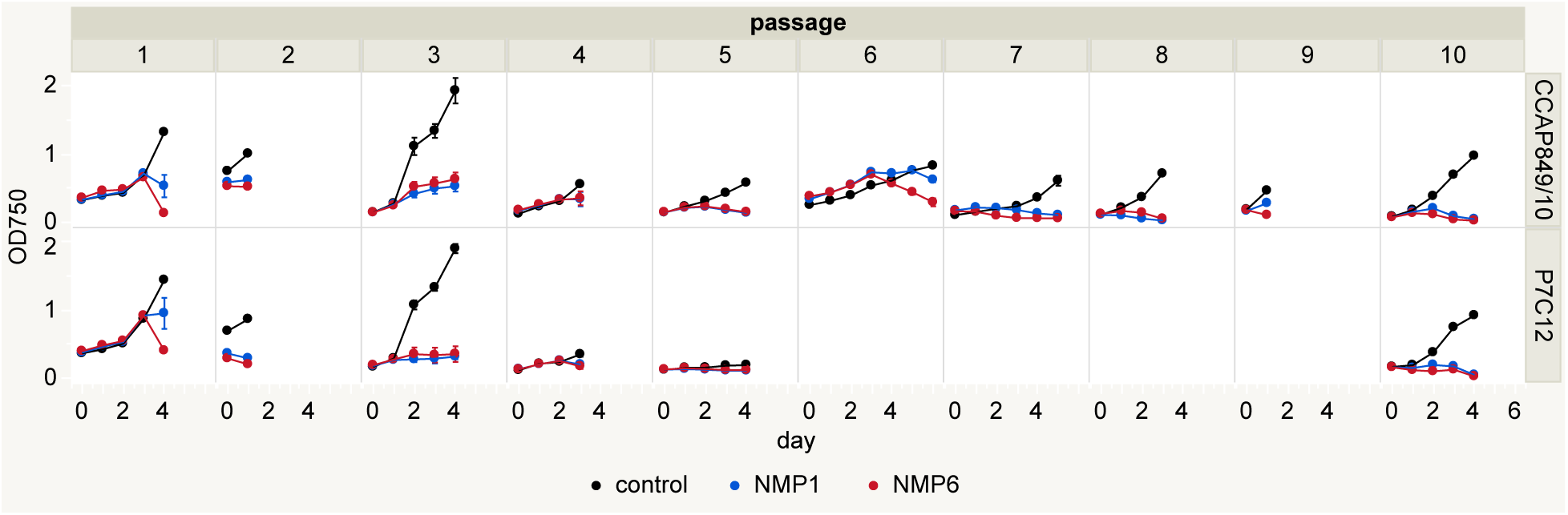
Optical density of 10 passages of two *N. oceanica* strains (CCAP 849/10 laboratory strain and P7C12 field-adapted strain) crossed with two bacterial infection sources NMP1 and NMP6.

**Table S1.**
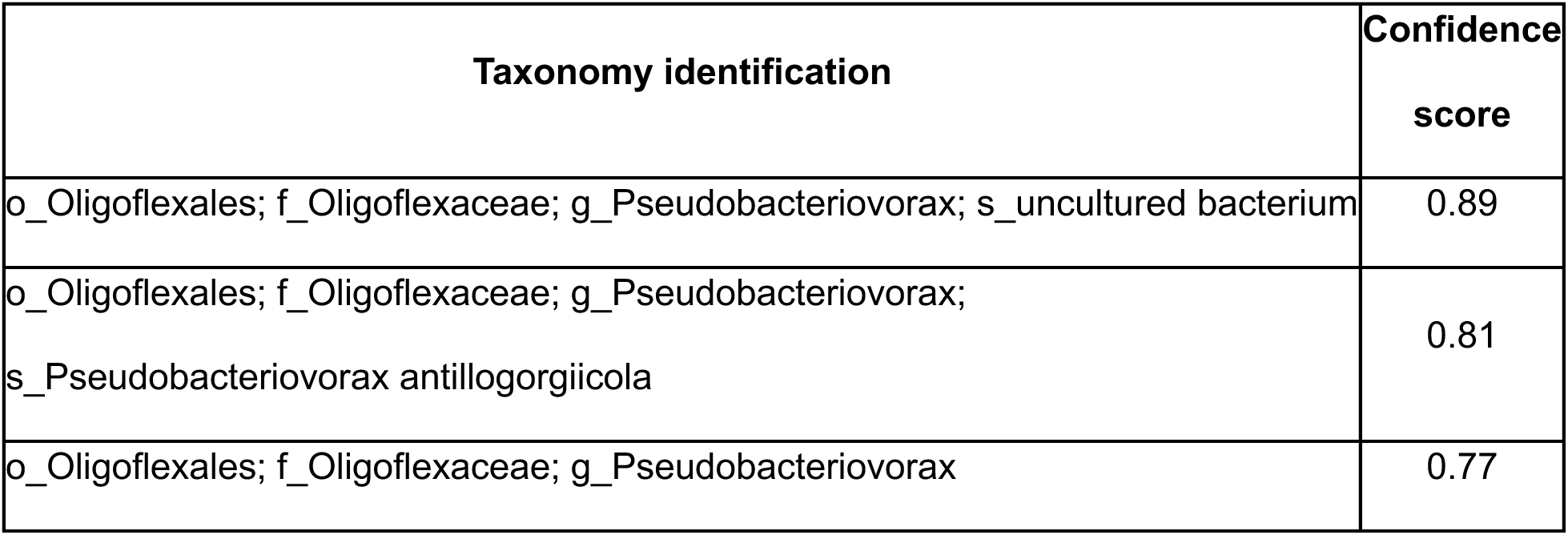
The sequencing reads of co-culture of *N. oceanica* and 2 infection sources were blasted again SILVA 138.2 SSURef NR99 full-length database focusing on the V3-V4 region of 16S rRNA gene. This analysis revealed the presence of three species in the Oligoflexaceae family, all belonging to the *Pseudobacteriovorax* genus. The only matching hit in the database was identified as *Pseudobacteriovorax antillogorgiicola* organism at the species level.

| Taxonomy identification | Confidence score |
| --- | --- |
| o_Oligoflexales; f_Oligoflexaceae; g_Pseudobacteriovorax; s_uncultured bacterium | 0.89 |
| o_Oligoflexales; f_Oligoflexaceae; g_Pseudobacteriovorax;<br>s_Pseudobacteriovorax antillogorgiicola | 0.81 |
| o_Oligoflexales; f_Oligoflexaceae; g_Pseudobacteriovorax | 0.77 |

**Table S2.** The absolute abundance of three different *Pseudobacteriovorax* species in the Oligoflexaceae family.

| Sample ID | *Pseudobacteriovorax*<br>sp. | *P. uncultured*<br>bacterium | *P.*<br>*antilogorgiicola* |
| --- | --- | --- | --- |
| CCAP84910_CONTROL_rep1_05202<br>024 | 0 | 0 | 0 |
| CCAP84910_CONTROL_rep1_05222<br>024 | 0 | 0 | 0 |
| CCAP84910_CONTROL_rep1_05242<br>024 | 0 | 0 | 0 |
| CCAP84910_CONTROL_rep2_05202<br>024 | 0 | 0 | 0 |
| CCAP84910_CONTROL_rep2_05222<br>024 | 0 | 0 | 0 |
| CCAP84910_CONTROL_rep2_05242<br>024 | 0 | 0 | 0 |
| CCAP84910_NMP1_rep1_05202024 | 207 | 135 | 0 |
| CCAP84910_NMP1_rep1_05222024 | 270 | 283 | 0 |
| CCAP84910_NMP1_rep1_05242024 | 143 | 42 | 0 |
| CCAP84910_NMP1_rep2_05202024 | 123 | 113 | 0 |
| CCAP84910_NMP1_rep2_05222024 | 426 | 256 | 0 |
| CCAP84910_NMP1_rep2_05242024 | 99 | 22 | 0 |
| CCAP84910_NMP6_rep1_05202024 | 271 | 226 | 0 |
| CCAP84910_NMP6_rep1_05222024 | 1432 | 718 | 2 |
| CCAP84910_NMP6_rep1_05242024 | 67 | 59 | 0 |
| CCAP84910_NMP6_rep2_05202024 | 480 | 235 | 0 |
| CCAP84910_NMP6_rep2_05222024 | 335 | 152 | 0 |
| CCAP84910_NMP6_rep2_05242024 | 65 | 13 | 0 |
| P7C12_CONTROL_rep1_05202024 | 0 | 0 | 0 |
| P7C12_CONTROL_rep1_05242024 | 0 | 0 | 0 |
| P7C12_NMP1_rep1_05202024 | 137 | 149 | 2 |
| P7C12_NMP1_rep1_05242024 | 0 | 0 | 0 |
| P7C12_NMP6_rep1_05202024 | 517 | 343 | 0 |
| P7C12_NMP6_rep1_05242024 | 66 | 29 | 0 |

**Table S3.** Number of forward and reserve sequence count (either forward or reverse direction) per each sample.

| Sample ID | Strain | Treatment | Replication | Day | Sequence Count |
| --- | --- | --- | --- | --- | --- |
| CCAP84910_CONTROL_rep1_05202024 | CCAP 849/10 | control | 1 | 0 | 174746 |
| CCAP84910_CONTROL_rep1_05222024 | CCAP 849/10 | control | 1 | 2 | 153121 |
| CCAP84910_CONTROL_rep1_05242024 | CCAP 849/10 | control | 1 | 4 | 52568 |
| CCAP84910_CONTROL_rep2_05202024 | CCAP 849/10 | control | 2 | 0 | 158844 |
| CCAP84910_CONTROL_rep2_05222024 | CCAP 849/10 | control | 2 | 2 | 220648 |
| CCAP84910_CONTROL_rep2_05242024 | CCAP 849/10 | control | 2 | 4 | 145189 |
| CCAP84910_NMP1_rep1_05202024 | CCAP 849/10 | NMP1 | 1 | 0 | 99007 |
| CCAP84910_NMP1_rep1_05222024 | CCAP 849/10 | NMP1 | 1 | 2 | 128376 |
| CCAP84910_NMP1_rep1_05242024 | CCAP 849/10 | NMP1 | 1 | 4 | 137647 |
| CCAP84910_NMP1_rep2_05202024 | CCAP 849/10 | NMP1 | 2 | 0 | 74679 |
| CCAP84910_NMP1_rep2_05222024 | CCAP 849/10 | NMP1 | 2 | 2 | 119673 |
| CCAP84910_NMP1_rep2_05242024 | CCAP<br>849/10 | NMP1 | 2 | 4 | 92075 |
| CCAP84910_NMP6_rep1_05202024 | CCAP<br>849/10 | NMP6 | 1 | 0 | 75157 |
| CCAP84910_NMP6_rep1_05222024 | CCAP<br>849/10 | NMP6 | 1 | 2 | 191248 |
| CCAP84910_NMP6_rep1_05242024 | CCAP<br>849/10 | NMP6 | 1 | 4 | 180795 |
| CCAP84910_NMP6_rep2_05202024 | CCAP<br>849/10 | NMP6 | 2 | 0 | 164038 |
| CCAP84910_NMP6_rep2_05222024 | CCAP<br>849/10 | NMP6 | 2 | 2 | 129807 |
| **CCAP84910_NMP6_rep2_05242024** | **CCAP**<br>**849/10** | **NMP6** | **2** | **4** | **38521** |
| P7C12_CONTROL_rep1_05202024 | P7C12 | control | 1 | 0 | 164409 |
| P7C12_CONTROL_rep1_05242024 | P7C12 | control | 1 | 4 | 72706 |
| P7C12_NMP1_rep1_05202024 | P7C12 | NMP1 | 1 | 0 | 129084 |
| P7C12_NMP1_rep1_05242024 | P7C12 | NMP1 | 1 | 4 | 108184 |
| P7C12_NMP6_rep1_05202024 | P7C12 | NMP6 | 1 | 0 | 149808 |
| P7C12_NMP6_rep1_05242024 | P7C12 | NMP6 | 1 | 4 | 89620 |

**Figure S2.**
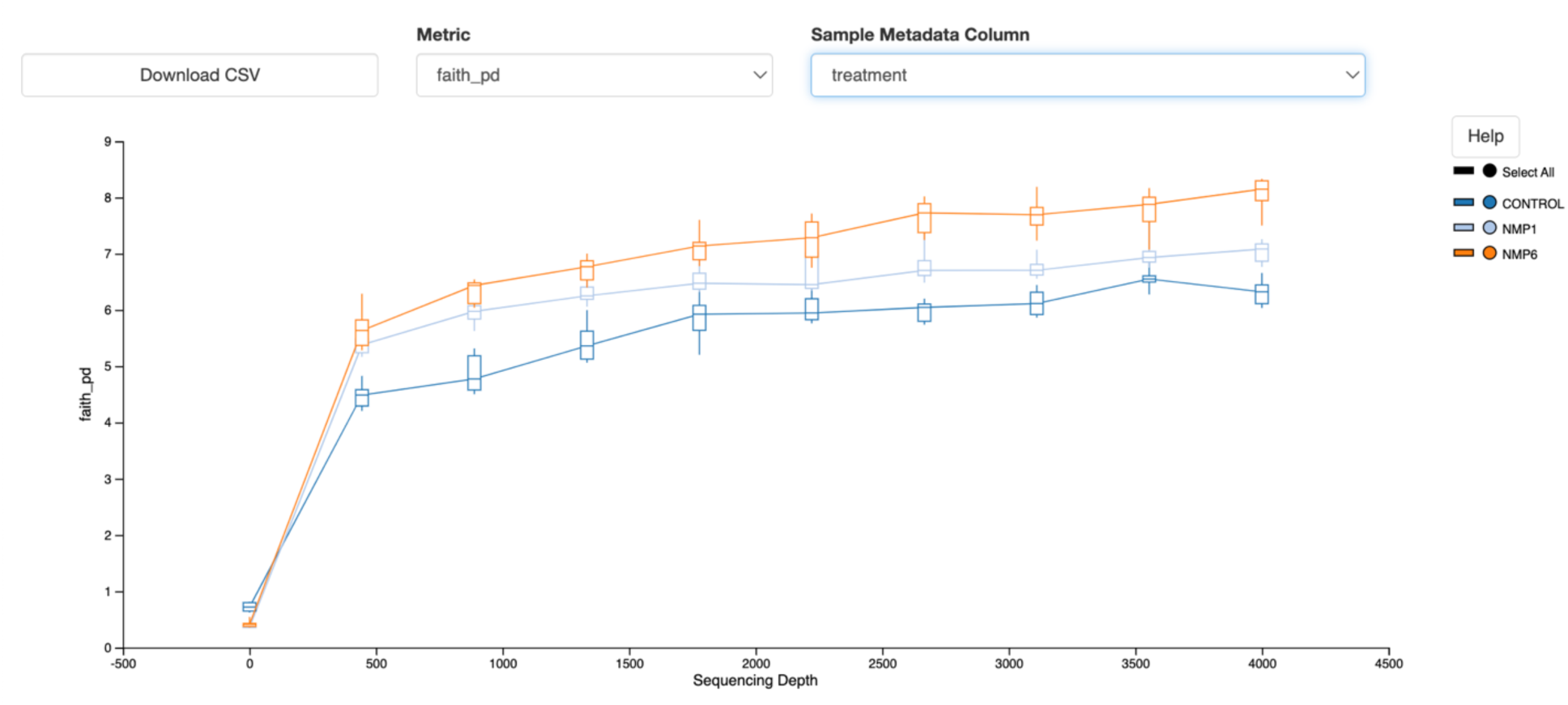
Alpha rarefaction plot by faith phylogenetic diversity (faith_pd) at multiple different sequencing depth. All the samples reach the stabilize point at around 4000 sequence count. All samples in this study were sequenced from ∼ 40,000 to ∼200,000 reads.

**Figure S3.**
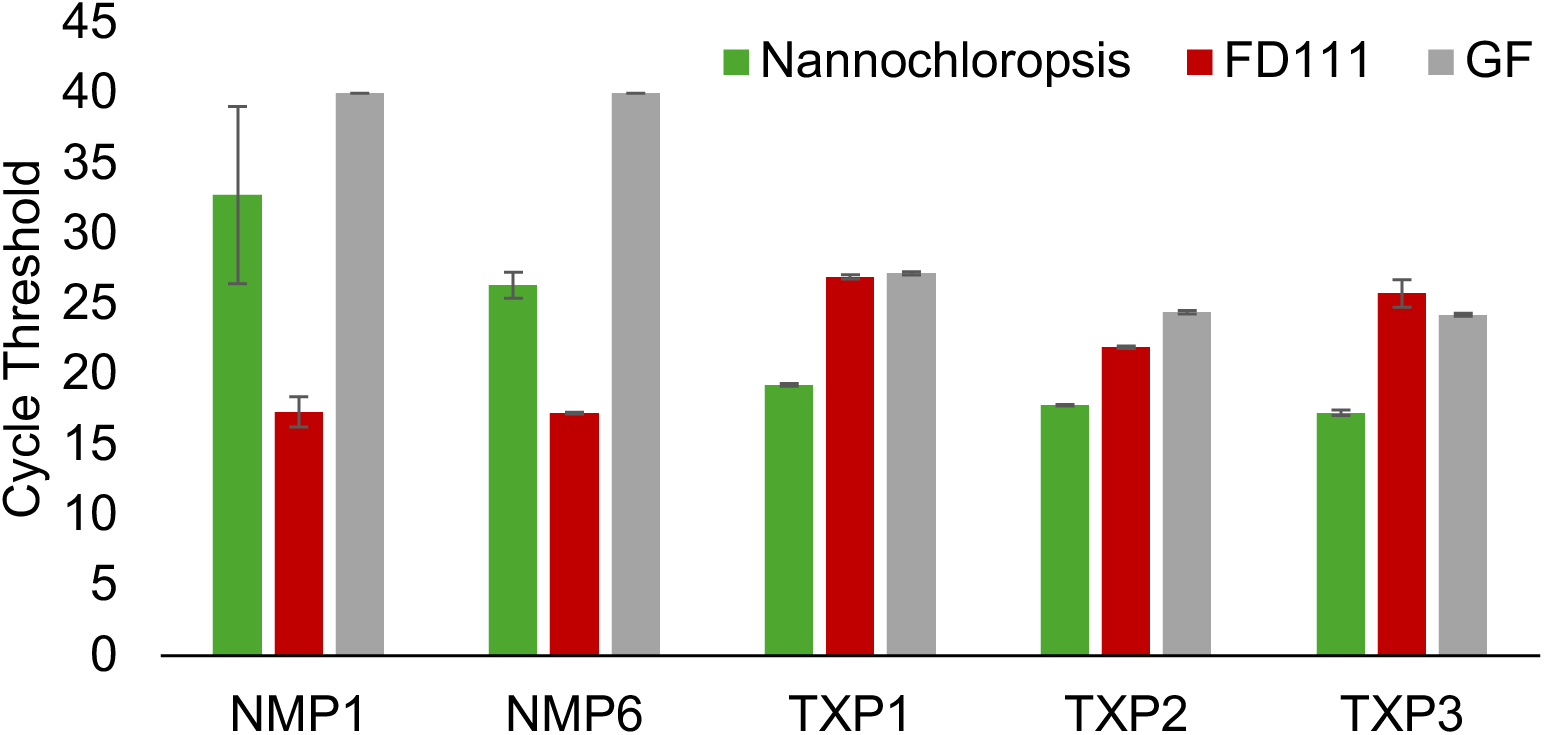
qPCR data from field samples taken from New Mexico Ponds NMP1 and NMP6, as well as Texas Ponds TXP1, 2, and 3. In NMP1 and NMP6, the FD111 bacterium was more prevalent compared to *Nannochloropsis* sp. and golden flagellate (GF), indicated by its lower cycle threshold. Conversely, in the TXP, the FD111 bacterium was less dominat, demonstrated by a higher cycle threshold value.

## Notes

### Competing Interest Statement

The authors have declared no competing interest.

## References

[1] K.M. Palanisamy, P. Bhuyar, M.A. Rahim, A. Vadiveloo, N.A. Al-Dhabi, N. Govindan, G. Maniam, Lipid enhancement in oleaginous Nannochloropsis sp. Under nitrate limitation for future bioenergy production, Int. J. Energy Res. 2023 (2023) 1–8.

[2] E.T. Chua, P.M. Schenk, A biorefinery for Nannochloropsis: Induction, harvesting, and extraction of EPA-rich oil and high-value protein, Bioresour. Technol. 244 (2017) 1416–1424.

[3] S. Kim, Y. Kwon, K.W. Kim, J.Y. Kim, Exploring the Potential of Nannochloropsis sp. Extract for Cosmeceutical Applications, Mar. Drugs 19 (2021). 10.3390/md19120690.

[4] G. Van Vooren, F. Le Grand, J. Legrand, S. Cuiné, G. Peltier, J. Pruvost, Investigation of fatty acids accumulation in Nannochloropsis oculata for biodiesel application, Bioresour. Technol. 124 (2012) 421–432.

[5] A. Taleb, J. Pruvost, J. Legrand, H. Marec, B. Le-Gouic, B. Mirabella, B. Legeret, S. Bouvet, G. Peltier, Y. Li-Beisson, S. Taha, H. Takache, Development and validation of a screening procedure of microalgae for biodiesel production: application to the genus of marine microalgae Nannochloropsis, Bioresour. Technol. 177 (2015) 224– 232.

[6] L. Novoveská, S.L. Nielsen, O.T. Eroldoğan, B.Z. Haznedaroglu, B. Rinkevich, S. Fazi, J. Robbens, M. Vasquez, H. Einarsson, Overview and challenges of large-scale cultivation of photosynthetic microalgae and Cyanobacteria, Mar. Drugs 21 (2023) 445.

[7] S.P. Fulbright, A. Robbins-Pianka, D. Berg-Lyons, R. Knight, K.F. Reardon, S.T. Chisholm, Bacterial community changes in an industrial algae production system, Algal Res. 31 (2018) 147–156.

[8] S. Bellou, G. Aggelis, Biochemical activities in Chlorella sp. and Nannochloropsis salina during lipid and sugar synthesis in a lab-scale open pond simulating reactor, J. Biotechnol. 164 (2012) 318–329.

[9] M. Kawaroe, J. Hwangbo, D. Augustine, H.A. Putra, Comparison of density, specific growth rate, biomass weight, and doubling time of microalgae Nannochloropsis sp. cultivated in Open Raceway Pond and Photobioreactor, Aacl Bioflux 8 (2015) 740– 750.

[10] P. Cunha, H. Pereira, M. Costa, J. Pereira, J.T. Silva, N. Fernandes, J. Varela, J.T. Silva, M. Simões, Nannochloropsis oceanica cultivation in pilot-scale raceway ponds—from design to cultivation, Appl. Sci. (Basel) (2020). 10.3390/app10051725.

[11] P. Aiyar, D. Schaeme, M. García-Altares, D. Carrasco Flores, H. Dathe, C. Hertweck, S. Sasso, M. Mittag, Antagonistic bacteria disrupt calcium homeostasis and immobilize algal cells, Nat. Commun. 8 (2017) 1756.

[12] S.A. Amin, L.R. Hmelo, H.M. van Tol, B.P. Durham, L.T. Carlson, K.R. Heal, R.L. Morales, C.T. Berthiaume, M.S. Parker, B. Djunaedi, A.E. Ingalls, M.R. Parsek, M.A. Moran, E.V. Armbrust, Interaction and signalling between a cosmopolitan phytoplankton and associated bacteria, Nature 522 (2015) 98–101.

[13] R.L. White, R.A. Ryan, Long-term cultivation of algae in open-raceway ponds: Lessons from the field, Ind. Biotechnol. (New Rochelle N. Y.) 11 (2015) 213–220.

[14] M.M. Rose, D. Scheer, Y. Hou, V.S. Hotter, A.J. Komor, P. Aiyar, K. Scherlach, F. Vergara, Q. Yan, J.E. Loper, T. Jakob, N.M. van Dam, C. Hertweck, M. Mittag, S. Sasso, The bacterium Pseudomonas protegens antagonizes the microalga Chlamydomonas reinhardtii using a blend of toxins, Environ. Microbiol. 23 (2021) 5525–5540.

[15] V. Hotter, D. Zopf, H.J. Kim, A. Silge, M. Schmitt, P. Aiyar, J. Fleck, C. Matthäus, J. Hniopek, Q. Yan, J. Loper, S. Sasso, C. Hertweck, J. Popp, M. Mittag, A polyyne toxin produced by an antagonistic bacterium blinds and lyses a Chlamydomonad alga, Proc. Natl. Acad. Sci. U. S. A. 118 (2021) e2107695118.

[16] B. Burgunter-Delamare, P. Shetty, T. Vuong, M. Mittag, Exchange or eliminate: The secrets of algal-bacterial relationships, Plants 13 (2024) 829.

[17] P.A. Lee, K.J.L. Martinez, P.M. Letcher, A.A. Corcoran, R.A. Ryan, A novel predatory bacterium infecting the eukaryotic alga Nannochloropsis, Algal Res. 32 (2018) 314–320.

[18] B. Humphrey, M. Mackenzie, M. Lobitz, J.Y. Schambach, G. Lasley, S. Kolker, B. Ricken, H. Bennett, K.P. Williams, C.R. Smallwood, J. Cahill, Biotic countermeasures that rescue Nannochloropsis gaditana from a Bacillus safensis infection, Front. Microbiol. 14 (2023) 1271836.

[19] S.A. Steichen, S. Gao, P. Waller, J.K. Brown, Association between algal productivity and phycosphere composition in an outdoor Chlorella sorokiniana reactor based on multiple longitudinal analyses, Microb. Biotechnol. 13 (2020) 1546–1561.

[20] S.A. Steichen, J.K. Brown, Real-time quantitative detection of Vampirovibrio chlorellavorus, an obligate bacterial pathogen of Chlorella sorokiniana, J. Appl. Phycol. 31 (2019) 1117–1129.

[21] R.M. Soo, B.J. Woodcroft, D.H. Parks, G.W. Tyson, P. Hugenholtz, Back from the dead; the curious tale of the predatory cyanobacterium Vampirovibrio chlorellavorus, PeerJ 3 (2015) e968.

[22] S.-H. Park, S.A. Steichen, X. Li, K. Ogden, J.K. Brown, Association of Vampirovibrio chlorellavorus with decline and death of Chlorella sorokiniana in outdoor reactors, J. Appl. Phycol. 31 (2019) 1131–1142.

[23] S. Attalah, P. Waller, S. Steichen, S. Gao, C.C. Brown, K. Ogden, J.K. Brown, Application of deoxygenation-aeration cycling to control the predatory bacterium Vampirovibrio chlorellavorus in Chlorella sorokiniana cultures, Algal Res. 39 (2019) 101427.

[24] B.J. Callahan, P.J. McMurdie, M.J. Rosen, A.W. Han, A.J.A. Johnson, S.P. Holmes, DADA2: High-resolution sample inference from Illumina amplicon data, Nat. Methods 13 (2016) 581–583.

[25] E. Bolyen, J.R. Rideout, M.R. Dillon, N.A. Bokulich, C.C. Abnet, G.A. Al-Ghalith, H. Alexander, E.J. Alm, M. Arumugam, F. Asnicar, Y. Bai, J.E. Bisanz, K. Bittinger, A. Brejnrod, C.J. Brislawn, C.T. Brown, B.J. Callahan, A.M. Caraballo-Rodríguez, J. Chase, E.K. Cope, R. Da Silva, C. Diener, P.C. Dorrestein, G.M. Douglas, D.M. Durall, C. Duvallet, C.F. Edwardson, M. Ernst, M. Estaki, J. Fouquier, J.M. Gauglitz, S.M. Gibbons, D.L. Gibson, A. Gonzalez, K. Gorlick, J. Guo, B. Hillmann, S. Holmes, H. Holste, C. Huttenhower, G.A. Huttley, S. Janssen, A.K. Jarmusch, L. Jiang, B.D. Kaehler, K.B. Kang, C.R. Keefe, P. Keim, S.T. Kelley, D. Knights, I. Koester, T. Kosciolek, J. Kreps, M.G.I. Langille, J. Lee, R. Ley, Y.-X. Liu, E. Loftfield, C. Lozupone, M. Maher, C. Marotz, B.D. Martin, D. McDonald, L.J. McIver, A.V. Melnik, J.L. Metcalf, S.C. Morgan, J.T. Morton, A.T. Naimey, J.A. Navas-Molina, L.F. Nothias, S.B. Orchanian, T. Pearson, S.L. Peoples, D. Petras, M.L. Preuss, E. Pruesse, L.B. Rasmussen, A. Rivers, M.S. Robeson 2nd, P. Rosenthal, N. Segata, M. Shaffer, A. Shiffer, R. Sinha, S.J. Song, J.R. Spear, A.D. Swafford, L.R. Thompson, P.J. Torres, P. Trinh, A. Tripathi, P.J. Turnbaugh, S. Ul-Hasan, J.J.J. van der Hooft, F. Vargas, Y. Vázquez-Baeza, E. Vogtmann, M. von Hippel, W. Walters, Y. Wan, M. Wang, J. Warren, K.C. Weber, C.H.D. Williamson, A.D. Willis, Z.Z. Xu, J.R. Zaneveld, Y. Zhang, Q. Zhu, R. Knight, J.G. Caporaso, Reproducible, interactive, scalable and extensible microbiome data science using QIIME 2, Nat. Biotechnol. 37 (2019) 852–857.

[26] E.P. McCauley, B. Haltli, R.G. Kerr, Description of Pseudobacteriovorax antillogorgiicola gen. nov., sp. nov., a bacterium isolated from the gorgonian octocoral Antillogorgia elisabethae, belonging to the family Pseudobacteriovoracaceae fam. nov., within the order Bdellovibrionales, Int. J. Syst. Evol. Microbiol. 65 (2015) 522–530.

[27] N. Kieffer, L. Poirel, C. Fournier, B. Haltli, R. Kerr, P. Nordmann, Characterization of PAN-1, a carbapenem-hydrolyzing class B β-lactamase from the environmental Gram-negative Pseudobacteriovorax antillogorgiicola, Front. Microbiol. 10 (2019) 1673.

[28] S.R. Starkenburg, A.A. Corcoran, Success through synergy (STS): Increasing cultivation yield and stability with rationally designed consortia, Los Alamos National Laboratory (LANL), Los Alamos, NM (United States); New Mexico Consortium, Los Alamos, NM (United States), 2022. 10.2172/1924073.

[29] M.R. Sanchez, E. Denning, T.C. Biondi, B. Hovde, S. Eacker, S. Getto, H. Kaur, A. Jebali, I. Echenique-Subiabre, M. Green, J. Gerber, B. Auch, F.O. Holguin, I. Liachko, H. Martinez, M. Balleza, J. Nalley, C. O’Kelly, J.B. Shurin, A.A. Corcoran, S.R. Starkenburg, Development of a field-deployable qPCR assay for real-time pest monitoring in algal cultivation systems, Algal Res. 74 (2023) 103194.

[30] S.F. Koval, S.H. Hynes, R.S. Flannagan, Z. Pasternak, Y. Davidov, E. Jurkevitch, Bdellovibrio exovorus sp. nov., a novel predator of Caulobacter crescentus, Int. J. Syst. Evol. Microbiol. 63 (2013) 146–151.

[31] R.J. Seidler, M.P. Starr, Isolation and characterization of host-independent Bdellovibrios, J. Bacteriol. 100 (1969) 769–785.

[32] A.R. Snyder, H.N. Williams, M.L. Baer, K.E. Walker, O.C. Stine, 16S rDNA sequence analysis of environmental Bdellovibrio-and-like organisms (BALO) reveals extensive diversity, Int. J. Syst. Evol. Microbiol. 52 (2002) 2089–2094.

[33] Y. Davidov, E. Jurkevitch, Diversity and evolution of Bdellovibrio-and-like organisms (BALOs), reclassification of Bacteriovorax starrii as Peredibacter starrii gen. nov., comb. nov., and description of the Bacteriovorax-Peredibacter clade as Bacteriovoracaceae fam. nov, Int. J. Syst. Evol. Microbiol. 54 (2004) 1439–1452.

